# The GENOMES UNCOUPLED1 protein has an ancient, highly conserved role in chloroplast gene expression but not in retrograde signalling

**DOI:** 10.1101/2022.02.25.481377

**Authors:** Suvi Honkanen, Ian Small

## Abstract

- The pentatricopeptide repeat protein GENOMES UNCOUPLED1 (GUN1) is required for chloroplast-to-nucleus signalling in response to plastid stress during chloroplast development in *Arabidopsis thaliana* but its exact molecular function remains unknown.
- We analysed GUN1 sequences in land plants and streptophyte algae. We tested functional conservation by complementation of the *Arabidopsis gun1* mutant with *GUN1* genes from the streptophyte alga *Coleochate orbicularis* or the liverwort *Marchantia polymorpha.* We also analysed the transcriptomes of *M. polymorpha gun1* knock-out mutant lines during chloroplast development.
- GUN1 evolved within the streptophyte algal ancestors of land plants and is highly conserved among land plants but missing from the *Rafflesiaceae* that lack chloroplast genomes. GUN1 genes from *C. orbicularis* and *M. polymorpha* restore chloroplast retrograde signalling and suppress the cold-sensitive phenotype of the *Arabidopsis gun1* mutant. However, GUN1 is not required for chloroplast retrograde signalling in the liverwort *M. polymorpha*.
- Our findings suggest that GUN1 is an ancient protein that evolved within the streptophyte algal ancestors of land plants before the first plants colonised land more than 470 million years ago. Its primary role is likely to be in chloroplast gene expression and its role in chloroplast retrograde signalling probably evolved more recently.

## Introduction

Plant chloroplasts evolved from endosymbiotic cyanobacteria. While chloroplasts retain their own genomes, the majority of the cyanobacterial genes were lost or have moved into the nuclear genome during evolution (Timmis *et al*., 2004). Consequently, many chloroplast protein complexes contain both nucleus- and chloroplast-encoded components (Rolland *et al*., 2012). Therefore, the expression of the chloroplast and nuclear genes needs to be tightly coordinated to ensure optimal chloroplast function during development and under changing environmental conditions (Chan *et al*., 2016). This coordination of chloroplast and nuclear gene expression requires both nucleus-to-chloroplast signals (anterograde signals) and chloroplast-to-nucleus signals (retrograde signals). Chloroplast-to-nucleus retrograde signalling mechanisms fall into two categories: biogenic retrograde signals that operate during early chloroplast development, and operational retrograde signals that are emitted by mature chloroplasts (reviewed in Pogson *et al*., 2008; Chan *et al*., 2016; Hernández-Verdeja & Strand, 2018; Wu & Bock, 2021).

Chloroplasts develop from small, undifferentiated plastids called proplastids. In *Arabidopsis* proplastid-to-chloroplast differentiation takes place in young cotyledons and in specific cells of the shoot apical meristem (Charuvi *et al*., 2012; Wu *et al*., 2018). Chloroplast stress during proplastid-to-chloroplast differentiation triggers biogenic chloroplast-to-nucleus signalling that results in strong transcriptional repression of <u>ph</u>otosynthesis-<u>a</u>ssociated <u>n</u>uclear <u>g</u>enes (phANGs), including genes encoding for light-harvesting chlorophyll a/b binding (LHCB) proteins and rubisco small sub-unit (RbcS) proteins (Harpster *et al*., 1984; Mayfield & Taylor, 1984).

While the exact identity of the chloroplast-emitted retrograde signal or signals is still being debated, the retrograde signalling mechanism that operates during proplastid-to-chloroplast differentiation in Arabidopsis involves the *GENOMES UNCOUPLED* (*GUN*) genes. The *genomes uncoupled* mutants were isolated based on their inability to repress transcription of the nuclear *LIGHT HARVESTING CHLOROPHYLL A/B-BINDING PROTEIN 1.2* (*LHCB1.2*) gene in response to plastid stress imposed by norflurazon treatment (Susek *et al*., 1993; Mochizuki *et al*., 2001; Woodson *et al*., 2011). Norflurazon inhibits carotenoid biosynthesis, which results in oxidative stress to the chloroplasts (Breitenbach *et al*., 2001). While five out of the six *GUN* genes (*GUN2*-*GUN6*) encode for proteins involved in tetrapyrrole metabolism, the molecular mechanism of *GUN1* function has remained unresolved. *GUN1* is unique among the other *GUN* genes in that it is also required for chloroplast-to-nucleus signalling when plastid gene expression is inhibited (Susek *et al*., 1993; Koussevitzky *et al*., 2007).

*GUN1* encodes for a pentatricopeptide peptide repeat (PPR) protein with a C-terminal small MutS-related (SMR) domain (Koussevitzky *et al*., 2007). PPR proteins constitute one of the largest protein families in seed plants, with 450 members in *Arabidopsis* (Barkan & Small, 2014). PPR proteins are characterised by tandem arrays of 31 to 36 amino-acid-long PPR motifs (Lurin *et al*., 2004). Most previously characterised PPR proteins function as sequence-specific RNA binding proteins. PPR proteins are encoded by the nuclear genome and almost exclusively localise to chloroplasts or mitochondria, where they bind their target RNAs. PPR proteins have been described to be involved in processing of their target RNAs in a variety of ways, including RNA cleavage, splicing, editing, stabilisation and translational activation (e.g. Pfalz *et al*., 2009; Prikryl *et al*., 2011; Wu *et al*., 2016; Aryamanesh *et al*., 2017; Zhou *et al*., 2017; Rojas *et al*., 2018; Lee *et al*., 2019a).

A subset of land plant PPR proteins, the PPR-SMR proteins, contain a C-terminal small MutS-related (SMR) domain. The SMR domain was first characterised in the MutS2 protein of the cyanobacterium *Synechocystis*, where it was reported to have DNA endonuclease activity (Moreira & Philippe, 1999; Fukui *et al*., 2007). SMR-domain-containing proteins are commonly found in both prokaryotic and eukaryotic organisms, although PPR proteins with SMR domains are (apart from a few exceptions) restricted to land plants and green algae (Liu *et al*., 2013). The *Arabidopsis* genome encodes for 8 PPR-SMR proteins, 5 of which locate to plastids. Three of these, PLASTID TRANSCRIPTIONALLY ACTIVE2 (pTAC2), SUPPRESSOR OF THYLAKOID FORMATION1 (SOT1) and SUPPRESSOR OF VARIEGATION7 (SVR7) are among the most abundant PPR proteins in plastids (Liu *et al*., 2013; Mergner *et al*., 2020). pTAC2 is an essential component of the plastid-encoded RNA polymerase complex and *ptac2* mutants are only able to grow on sucrose-supplemented media (Pfalz *et al*., 2006). SOT1 facilitates proper assembly and maturation of the plastid ribosome by processing the plastid 23S-4.5S ribosomal RNA (rRNA) precursor (Wu *et al*., 2016; Zhou *et al*., 2017). *svr7* mutants also possess plastid rRNA processing and gene expression defects, most notably impaired expression of the plastid ATP synthase (Liu *et al*., 2010; Zoschke *et al*., 2013). Therefore, PPR-SMR proteins are required for appropriate plastid gene expression.

In contrast to the other chloroplast-localised Arabidopsis PPR-SMR proteins, the GUN1 protein only accumulates in specific tissues (Mergner *et al*., 2020). While the GUN1 transcript is abundantly expressed, GUN1 protein accumulation coincides with proplastid-to-chloroplast differentiation in the cotyledons during early seedling development and in specific cells of the shoot apical meristem, based on GUN1-green fluorescent protein (GFP) fusion signal (Wu *et al*., 2018). In addition, the GUN1-GFP protein accumulates in response to inhibitors that induce plastid retrograde signalling (Wu *et al*., 2018). Several recent studies have proposed roles for the *Arabidopsis* GUN1 protein based on its interactions with other chloroplast-localised proteins. These include interaction with the chaperone cpHSC70-1 to control chloroplast protein import (Wu *et al*., 2019; Tadini *et al*., 2020), interaction with nucleus-encoded plastid RNA polymerase (NEP) to control the accumulation of NEP-encoded transcripts (Tadini *et al*., 2020), interaction with multiple tetrapyrrole biosynthesis enzymes, heme and porphyrins to control the flux through the tetrapyrrole biosynthesis pathway (Shimizu *et al*., 2019), and interaction with MULTIPLE ORGANELLAR RNA EDITING FACTOR2 (MORF2) to control RNA editing (Zhao *et al*., 2019).

The numerous proposed GUN1 interaction partners and mechanisms of function are intriguing but confusing. Current data on GUN1 function is limited to *Arabidopsis*, so we set out to investigate the origin and evolution of the land plant GUN1 proteins. Here we identify GUN1 as an ancient protein that is highly conserved across land plants. The pattern of amino acid conservation along the GUN1 protein is consistent with the hypothesis that GUN1, like other characterised PPR proteins, encodes for a nucleic acid binding protein. Finally, the retrograde signalling cascade that functions downstream of the conserved core GUN1 mechanism is not conserved and may have evolved more recently.

## Materials and Methods

### Phylogenetic analysis

*Arabidopsis* and *Marchantia* GUN1 sequences were retrieved from TAIR (https://www.arabidopsis.org/) and MarpoIBase (https://marchantia.info/), respectively. Full-length GUN1 sequences were obtained from a representative set of land plants by BLAST searches using the *Arabidopsis* sequence to search GenBank and selected translated sequence sets (whole genome shotgun or transcriptome shotgun assemblies) via the NCBI Sequence Set Browser (https://www.ncbi.nlm.nih.gov/Traces/wgs/). A set of 76 phylogenetically diverse GUN1 sequences (including representatives from algae, bryophytes, lycophytes, ferns, gymnosperms and angiosperms) were aligned using the G-INS-i algorithm in MAFFT v7 (Katoh & Standley, 2013). The most highly conserved region of this alignment (876 positions) was used to generate a GUN1 sequence profile with *hmmbuild* from the HMMER package (v3.3.1) (http://hmmer.org; Eddy, 2011), which in turn was used to search for GUN1 sequences (using *hmmsearch* with default parameters) in translations of various transcriptome datasets, most notably putative PPR protein sequences compiled by Gutmann *et al*. (2020) from the 1KP data set (Carpenter *et al*., 2019; Leebens-Mack *et al*., 2019). The 1KP transcriptomes were filtered to remove those encoding fewer than 10000 distinct proteins to avoid trivial false negatives due to low coverage and those from organisms other than green algae and land plants. This resulted in 1128 analysable samples from 894 plant species. Specific searches were also made in data sets of particular interest, or where GUN1 could not be found in the corresponding 1KP samples. These additional data sets included whole genome shotgun data from *Sapria himalayana* (Cai *et al*., 2021) and whole transcriptome data from *Rafflesia cantleyi* (Lee *et al*., 2016), both holo-parasites from the *Rafflesiaceae*.

### Plant lines and growth conditions

*Arabidopsis thaliana* wild type Columbia (Col-0) and *Atgun1* T-DNA insertion line SAIL_ 290_D09 were used in this study. Arabidopsis seeds were sterilised with chlorine gas for 2-4 hours and grown on sterile ½ MS medium (PhytoTech Labs) under long-day conditions (16 hours light 8 hours dark) at 100 μmol.m^−2^.sec^−1^ at 22 °C unless otherwise mentioned.

*M. polymorpha* wild-type accessions Takaragaike-1 (Tak-1) male and Tagarakaike-2 (Tak-2) female were kindly provided by Prof. John Bowman, Monash University. *M. polymorpha* CRISPR knock-out lines *Mpgun1-1* and *Mpgun1-2* were generated in this study. *M. polymorpha* plants were grown on sterile ½ Gamborg’s medium (Duchefa Biochemie) supplemented with 1.2 % agar under long day conditions (16 hours light 8 hours dark) unless otherwise mentioned. Crossing and spore sterilisation were carried out as described in methods S1.

### Generation of *A. thaliana gun1* mutant complementation lines

All primer sequences are provided in Table S1. Annotated sequences of the complementation constructs are provided in Notes S1. AtGUN1, MpGUN1 and CoGUN1 protein coding sequences were aligned using Geneious (Kearse *et al*., 2012). The region encoding for chloroplast transit peptides was estimated based on TargetP-2.0 transit peptide cleavage site predictions (Almagro Armenteros *et al*., 2019) and lack of conservation between the protein sequences. The *MpGUN1* coding sequence excluding the putative chloroplast transit peptide encoding region (amino acids 1-114) was amplified from *M. polymorpha* cDNA using PrimeSTAR HS DNA polymerase (Takara Bio) with primers CDS_MpGUN1_At_gib_F and CDS_MpGUN1_At_gib_R. The *CoGUN1* coding sequence excluding the chloroplast transit peptide encoding region (amino acids 1-145) was ordered as gene synthesis fragment from Integrated DNA Technologies (IDT). A fragment containing the *AtGUN1* promoter, 5’ untranslated region and chloroplast transit peptide (amino acids 1-135) was amplified from Arabidopsis Col-0 genomic DNA with primers pro_AtGUN1_gib_F and tp_AtGUN1_Mp_gib_R for the MpGUN1 construct, and primers pro_AtGUN1_gib_F and tp_AtGUN1_Co_gib_R for the CoGUN1 construct. The AtGUN1 terminator fragment was amplified with primers term_AtGUN1_Mp_gib_F and term_AtGUN1_gib_R for the MpGUN1 construct and with primers term_AtGUN1_Co_gib_F and term_AtGUN1_gib_R for the CoGUN1 construct. The three fragments for each construct were assembled on a *XbaI-* and-*SacI*-digested gel-purified MpGWB101 vector (Ishizaki *et al*., 2015) using Gibson assembly. To create a control construct for the complementation of the *Atgun1* mutant with *AtGUN1* genomic fragment the wild type *AtGUN1* gene sequence was amplified from Col-0 DNA using primers pro_AtGUN1_gib_F and term_AtGUN1_gib_R and cloned onto the *XbaI*-and-*SacI*-linearised MpGWB101 vector backbone.

*A. thaliana* wild-type and *gun1* mutant plants were transformed using the floral dip method (Zhang *et al*., 2006) with binary vectors described above using *Agrobacterium* strain GV3101. Seeds were surface-sterilised and sown on sterile plates supplemented with 15 μg⋅ ml^−1^ hygromycin. After one week, hygromycin-resistant plants were transferred on soil. The genotype of *Arabidopsis* wild type, *gun1* and *gun1* complemented with *AtGUN1*, *MpGUN1* and *CoGUN1* was PCR-verified using the Phire plant direct PCR kit (Thermo Fisher) as recommended by the manufacturer. The presence of the T-DNA insertion was verified using primers LBb1.3 and SAIL_280_D09_LP, whereas the presence/absence of intact wild-type *GUN1* sequence was verified with primers SAIL_280_D09_LP and SAIL_280_D09_RP. Seeds from 10–15 independent transformant lines were observed for the initial phenotypic assessment. All final phenotype and gene expression analyses were carried out on non-segregating T2 or T3 seeds of three independent homozygous transformant lines for each construct.

### *M. polymorpha* transformation and generation of transgenic *Mpgun1* CRISPR/Cas9 knock-out lines

*M. polymorpha* transformation and generation or transgenic *Mpgun1* CRISPR/Cas9 knock-out lines is described in supporting methods S1-S2.

### Plant growth experiments

#### Lincomycin treatment of *A. thaliana*

Arabidopsis seeds were sterilised and plated on sterile ½ MS medium pH 5.7, 2 % sucrose, 0.8 % agar. For the lincomycin treatment the growth media was supplemented with 220 μg⋅ml^−1^ lincomycin (Sigma). The seeds were stratified at 4 °C for 2 days, after which the plants were grown at 22 °C under continuous light at 100 μmol⋅m^−2^⋅s^−1^ for 5 days before imaging or harvesting the tissue for RNA extraction.

#### Combined low light and lincomycin treatment of *A. thaliana*

Arabidopsis seeds were sterilised and plated on sterile ½ MS medium pH 5.7 without sucrose, 0.8 % agar. For the lincomycin treatment the growth media was supplemented with 220 μg⋅ml^−1^ lincomycin (Sigma). The seeds were stratified at 4 °C for 4 days, and then grown at 22 °C under continuous light at 1 μmol⋅m^−2^⋅s^−1^ for 3 days before imaging.

#### *A. thaliana* cold growth experiment

Arabidopsis seeds were sterilised and plated on ½ MS medium pH 5.7 without sucrose, 0.8 % agar. The plates were placed directly at 4 °C growth room under long day conditions 8 hours dark 16 hours light at 100 μmol⋅m^−2^⋅s^−1^. Plants were imaged after 7 weeks.

#### Spectinomycin and norflurazon treatments of *M. polymorpha* spores

*M. polymorpha* spores were sterilised and plated on ½ Gamborg’s medium (Duchefa Biochemie) supplemented with 1.2 % agar and 500 μg⋅ml^−1^ spectinomycin or 5 μM norflurazon. The plates were placed under long day conditions for 48 hours, after which the spores were resuspended in 1 ml of sterile water, transferred into a microcentrifuge tube and spun down at 6,000 rpm for 1 minute. The water was removed, and the spore pellet flash-frozen in liquid nitrogen.

#### RNA extraction

Extraction of total RNA from *Arabidopsis* seedlings and *Marchantia* spores was carried out using the Direct-Zol RNA MINIprep kit (Zymo Research) following the manufacturer’s protocol. Three independent biological replicates were extracted for each line. RNA was quantified using a NanoDrop spectrophotometer (Thermo Fisher) and diluted to 250 ng⋅μl^−1^ before DNase treatment using Turbo DNase (Ambion) as recommended by the manufacturer.

#### cDNA synthesis and quantitative real time PCR (qPCR)

One μg of DNase-treated RNA was used as a template for cDNA synthesis. cDNA was generated with oligo dT18 primer using the Protoscript II reverse transcriptase (NEB) in the presence of Murine RNase inhibitor (NEB). cDNA was diluted 1:5 prior to qPCR. qPCR was carried out using SYBR Premix Ex Taq II (Takara Bio) qPCR reagent. Each 5 μl reaction contained 1 μl primer mix, 1.5 μl cDNA dilution and 2.5 μl Sybr II master mix. Each biological replicate sample was run in three technical replicates. The amplification was carried out in a LightCycler480 instrument (Roche Diagnostics) using the following cycling conditions: initial denaturation 1 min at 95, then 40 cycles 10 s at 95 °C, 10 s at 60 °C and 20 s at 72 °C. For primer sequences see Table S1. The data was analysed using LinRegPCR (Ruijter *et al*., 2009) version 2017.1. The expression of each gene of interest was first separately normalised against the reference genes *AtPDF2* and *AtUBQ10* for *Arabidopsis* and *MpEF1α* and *MpACT* for *M. polymorpha*. The geometric mean of the two normalised values was then recorded as the expression level.

#### RNA sequencing

Transcriptome libraries of *M. polymorpha* wild-type and *Mpgun1* mutant spores were prepared using 200 ng of DNase-treated total RNA as a template for the TruSeq Stranded Total RNA kit with Ribo-Zero Plant (Illumina). The libraries were sequenced on an Illumina HiSeq 4000 platform (150 nt paired-end reads) at Novogene, Hong Kong. At least 5.7 GB raw data was obtained for each replicate library. Sequencing read data was deposited at the Short Read Archive database at the National Center for Biotechnology Information under project number PRJNA800059.

Optical duplicate reads were first removed with clumpify (parameters: dedupe optical dist = 40) from the bbmap package (https://sourceforge.net/projects/bbmap/) and adapters were trimmed with bbduk (parameters: ktrim=r k=23 mink=11 hdist=1 tpe tbo ftm=5). The reads were then assigned to transcripts using Salmon v1.3.0 (Patro *et al*., 2017) (parameters: −1 A -- validateMappings) against an index prepared with the *M. polymorpha* MpTak_v6.1 reference genome and cDNA assemblies (https://marchantia.info/). Differential expression analyses were carried out using DESeq2 (Love *et al*., 2014). Functional annotations for MpTak_v5.1 genome release were used to annotate differentially expressed genes (log_2_ fold-change >1 or <-1 and padj < 0.01) and to identify *M. polymorpha* photosynthesis-associated nuclear genes. Gene Ontology (GO) enrichment analyses were performed on the Dicots Plaza 4.5 platform (Van Bel *et al*., 2018) using standard settings with differentially expressed genes showing log2 fold-change >1 or <−1 and padj < 0.01 as an input.

#### Microscopy and image analysis

Arabidopsis seedlings and *M. polymorpha* gemmae were imaged using an Olympus Camedia C 7070 camera mounted on a Leica SZ61 dissecting microscope or with an Apple iPad (7^th^ generation). Transmitted light microscopy images of *M. polymorpha* spores were obtained using an Olympus Camedia C 7070 camera mounted on a Olympus BX51 microscope. Image analysis to quantify phenotypical differences was carried out using Fiji (Schindelin *et al*., 2012). Images were adjusted using Adobe Photoshop 2020.

## Results

### GUN1 evolved within the streptophyte algae and is conserved among land plants

To discover when GUN1 arose and its distribution within the plant lineage, we searched for GUN1-like sequences using the *Arabidopsis* GUN1 (AtGUN1) sequence as a query in *blastp* (Altschul *et al*., 1990) searches of the GenBank non-redundant protein database. Likely GUN1 sequences were verified by alignment to AtGUN1, by reciprocal *blastp* to Arabidopsis protein sequences and by identification of the expected PPR motifs and C-terminal SMR domain. Land plants form a monophyletic group that evolved from the streptophyte algae (Wickett *et al*., 2014). We identified putative GUN1 sequences in the streptophyte algal groups most closely related to land plants (*Zygnematales*, *Coleochaetales* and *Charales*), but not in other streptophyte algal groups (*Klebsormidiales*, *Mesostigmatales* and *Chlorokybales*) nor in other green algae (*Chlamydomonadales*) (Figure S1) This suggests the GUN1 protein is ancient and evolved in the streptophyte algal ancestors of land plants before the first plants colonised land.

To systematically assess the conservation of GUN1 among plants, we selected 76 phylogenetically diverse full-length GUN1 sequences from a representative set of land plants and streptophyte algae (Figure 1, Figure S1), aligned them and retained the most conserved region of the alignment (which included all of the PPR motifs and the SMR domain). From this alignment we developed a sequence profile using *hmmbuild* (http://hmmer.org; Eddy, 2011) and used it to screen ~500,000 PPR sequences derived from the 1KP plant transcriptome data set (Carpenter *et al*., 2019; Leebens-Mack *et al*., 2019; Gutmann *et al*., 2020) with *hmmsearch*. Putative GUN1 orthologues could be distinguished from other PPR-SMR sequences based on the domain score reported for each match by *hmmsearch* (Figure S2, Table S2). *GUN1* transcripts were detected in 824 species out of the 894 analysed. We identified conserved GUN1 sequences in 345 of the 366 plant families represented in this dataset. Notably, we found that transcriptomes of non-photosynthetic parasitic plants with highly reduced plastid genomes, such as *Pilostyles thunbergii* (Bellot & Renner, 2016), *Balanophora fungosa* (Su *et al*., 2019) and *Conopholis americana* (Wicke *et al*., 2013) encode GUN1. Of the 21 families apparently lacking GUN1, 14 are green algal families outside the streptophyte clades closely related to land plants, and thus not expected to contain GUN1. One of the remaining seven is represented in the original set of 76 sequences used to build the GUN1 profile, thus does contain GUN1, even though no GUN1 transcripts were detected in the 1KP sample. This left six land plant families expected to contain GUN1 (*Agapanthaceae, Cyrillaceae, Hymenophyllaceae, Juglandaceae, Monocleaceae, Thelypteridaceae*) for which the data so far did not provide clear evidence of GUN1 sequences (Table S2). After searching in additional whole transcriptome or whole genome shotgun sequencing datasets, we were able to identify putative GUN1 sequences from *Agapanthus*, *Hymenophyllum* and *Juglans*. For the three remaining families, each represented by only a single sample in our 1KP dataset, we were unable to find additional data to search. We also searched whole transcriptome shotgun sequences from *Rafflesia cantleyi* (Lee *et al*., 2016) and whole genome shotgun sequences from *Sapria himalayana* (Cai *et al*., 2021) (both *Rafflesiaceae*). These two closely related genera are claimed to lack a plastid genome. We found no putative GUN1 sequences in either dataset. *Sapria himalayana* is the only embryophyte for which we were unable to find a putative *GUN1* gene in its genome, and *Rafflesia cantleyi* one of the few for which we were unable to find putative *GUN1* transcripts in its transcriptome.

**Fig. 1.**
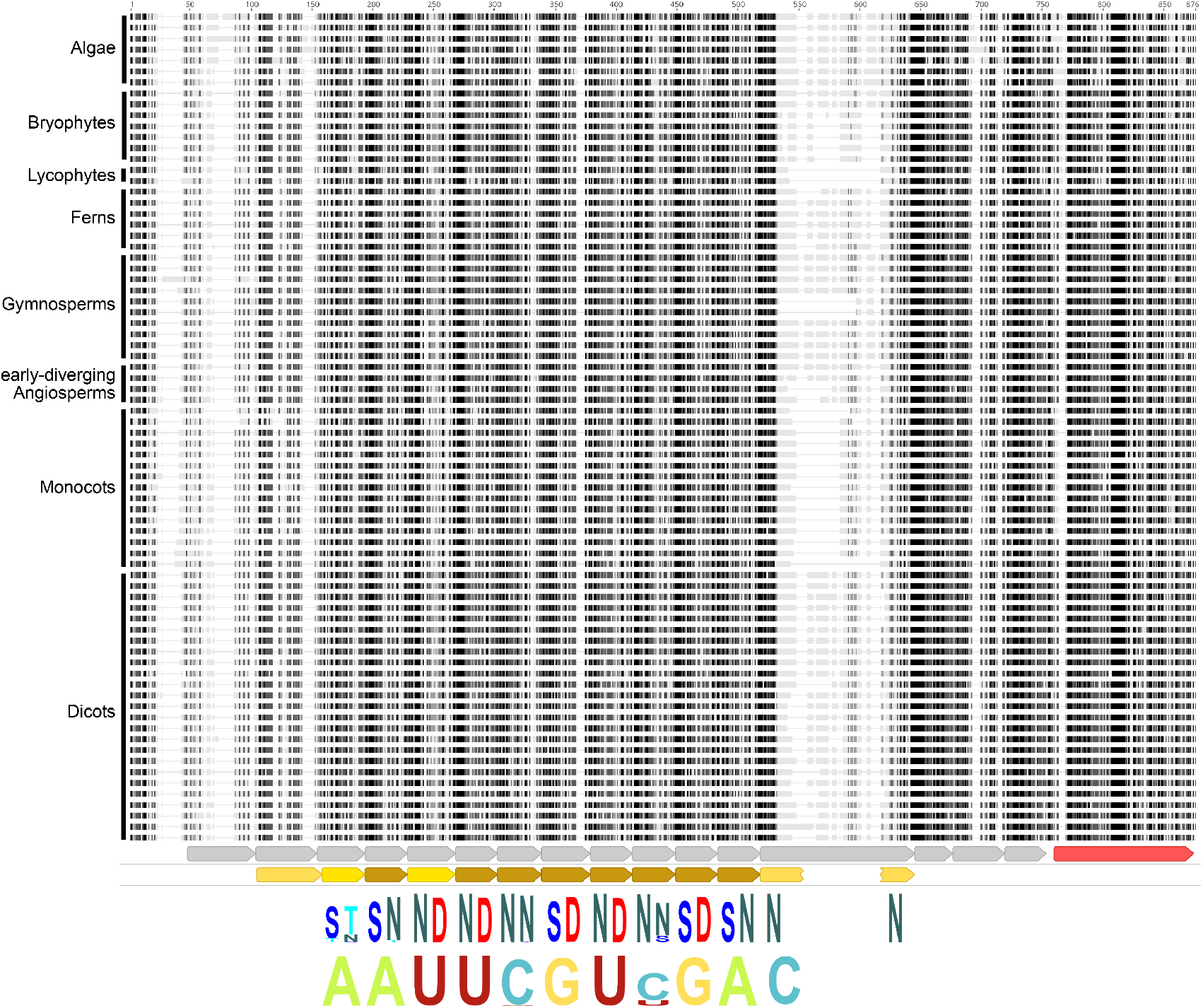
*GUN1* is conserved in most streptophytes. The top part of the figure represents a multiple alignment of 76 GUN1 protein sequences from diverse streptophyte algae and land plants. The alignment was constructed using MAFFT and visualised with Geneious. Only the central, most conserved region of the alignment is shown (corresponding to positions 135–888 in AtGUN1). Darker shading indicates higher similarity to the consensus. The sequences are grouped (approximately) by phylogenetic relationship as indicated; for the species names and amino acid sequences, see the alignment in Figure S1. Below the alignment, the annotation tracks show helix-turn-helix motifs (grey arrows) and SMR domain (red arrow) predicted by Alphafold and PPR motifs (brown arrows) predicted by hmmsearch using a P-type PPR motif HMM. Dark brown motifs have higher confidence. The final PPR motif is interrupted by a long insertion but Alphafold predicts that the N- and C-terminal helices nevertheless interact together as in a typical PPR motif. Below these motif predictions, the fifth and last amino acids of the conserved PPR motifs are indicated as sequence logos generated by Skylign (Wheeler *et al*., 2014) (letter height represents information content) from the alignment of 893 GUN1 sequences (the 76 sequences shown here plus 817 sequences from the 1KP PPR dataset). Finally, at the bottom of the figure, the expected RNA binding specificities of the conserved PPR motifs are indicated, again as a Skylign sequence logo. The HMM profile and full alignment of all 893 identified GUN1 sequences are obtainable from Dryad (https://doi.org/10.5061/dryad.x0k6djhmk).

### The predicted binding specificity of GUN1 proteins is conserved

Conservation of protein function is likely to depend on the level of sequence conservation in the functionally important regions of the protein. The GUN1 protein consists of 12-16 pentatricopeptide repeat (PPR) motifs (depending on the approach used to recognise and define the motifs) and a C-terminal small MutS-related (SMR) domain. Figure 1 shows the alignment of the conserved regions of the GUN1 sequences used to generate the HMM profile, and the structural motifs that correspond, as predicted by Alphafold (Jumper *et al*., 2021; Varadi *et al*., 2022) or hmmsearch (Eddy, 2011). Alphafold predicts 16 helix-turn-helix motifs of which 12 are recognised as PPR motifs by hmmsearch with a P-type PPR HMM (Cheng *et al*., 2016). The first helix-turn-helix motif predicted by Alphafold is poorly conserved between species, but the final three are highly conserved within GUN1 sequences although quite divergent in sequence and length from typical PPR motifs.

The binding specificity of PPR proteins can be predicted based on the 5^th^ and last amino acid of each PPR motif (Barkan *et al*., 2012). The amino acids at these positions of PPR motifs 2– 12 are highly conserved between all land plant and streptophyte algae GUN1 protein sequences (Figure 1, Figure S1). When these amino acid combinations are used to predict the nucleotide most likely to be bound by each motif, this conservation is even more striking, with the only predicted variation being at motifs 6 and 9 where different combinations are predicted to bind either C or U. This may not actually reflect functional divergence, as any of the combinations are likely to have high affinity for both C and U (Yan *et al*., 2019). The structure predicted by Alphafold forms a contiguous solenoid similar to that of other PPR proteins that are known to bind RNA, albeit with some idiosyncracies. The conservation of the PPR motifs, and in particular the residues that in other PPR proteins define RNA binding specificity, is consistent with the hypothesis that GUN1 encodes for an RNA binding protein with a highly conserved target. Nevertheless, the predicted target sequence in Figure 1 is unlikely to be correct; this 11-nucleotide sequence occurs in many chloroplast genomes but not all, and not at conserved positions.

### GUN1 SMR domains are conserved

The GUN1 protein contains a C-terminal SMR domain that is highly conserved between all the full-length GUN1 sequences in our dataset. Four SMR-domain-containing proteins have been reported to have endonuclease activity in other organisms (Liu *et al*., 2013). Subfamily 2 SMR proteins that have endonuclease cleavage activity have two conserved motifs: the LDXH motif in the N-terminus of the SMR domain and a central TGXG motif (Watanabe *et al*., 2003; Bhandari *et al*., 2011). The central TGXG motif is perfectly conserved as TGWG in all full-length GUN1 proteins in our data set. The N-terminal LDXH motif is also almost perfectly conserved as LDLH, with a scattering of exceptions that have VDLH in this position. These observations show that the SMR domains of land plant GUN1 proteins are conserved and most likely contribute to GUN1 function.

### *GUN1* genes from the streptophyte alga *C. orbicularis* and the liverwort *M. polymorpha* restore chloroplast retrograde signalling in the *Atgun1* mutant

As the predicted binding specificity of GUN1 proteins is conserved among land plants and in the streptophyte algae, we hypothesised that GUN1 proteins are also functionally conserved. To test this hypothesis, we assessed whether *GUN1* genes from the streptophyte alga *Coleochaete orbicularis* and the liverwort *Marchantia polymorpha* can complement the Arabidopsis *gun1* knock-out mutant. *C. orbicularis* and *M. polymorpha* each contain a single-copy *GUN1* gene. The *C. orbicularis* GUN1 protein shares 42.5 % amino acid identity (57.2 % match using BLSM62) with AtGUN1, while the *M. polymorpha* GUN1 protein is 48.4 % identical (66.4 % match using BLSM62) with AtGUN1. Sequences corresponding to chloroplast transit peptides in CoGUN1 and MpGUN1 were estimated based on lack of sequence conservation and replaced with the *Arabidopsis* GUN1 transit peptide. These coding sequences were then expressed from the *AtGUN1* promoter in the *Atgun1* (SAIL_ 290_D09) mutant background. As a control the *Atgun1* mutant was complemented with an *AtGUN1* genomic fragment.

In wild-type *Arabidopsis*, inhibition of plastid translation during chloroplast development activates chloroplast retrograde signalling that results in strong transcriptional repression of photosynthesis-associated nuclear genes (phANGs). In the *Atgun1* mutant this transcriptional repression is defective, although not completely abolished (Figure 2, Koussevitzky *et al*., 2007). We hypothesised that *CoGUN1* and *MpGUN1* can restore the GUN1-dependent repression of phANG in the *Atgun1* mutant when plastid translation is inhibited. To test this hypothesis, we germinated the *Atgun1* seeds expressing *CoGUN1*, *MpGUN1* or *AtGUN1* on media supplemented with 200 μg⋅ml^−1^ lincomycin, an inhibitor of plastid translation. As expected, the inhibition of plastid translation by lincomycin resulted in strong reduction in steady-state transcript levels of four phANGs, i.e. *AtLHCB1.2*, *AtLHCB2.2*, *AtCA1* and *AtACP12* in wild-type *Arabidopsis*, but less so in the *Atgun1* mutant (Figure 2). Expression of *CoGUN1* and *MpGUN1* in the *Atgun1* mutant background restored the repression of these four phANG transcripts to wild-type levels (Figure 2). These results indicate *CoGUN1* and *MpGUN1* can activate the *AtGUN1*-dependent retrograde signalling cascade that leads to transcriptional repression of photosynthesis-associated nuclear encoded genes (phANGs) in *Arabidopsis*.

**Fig. 2.**
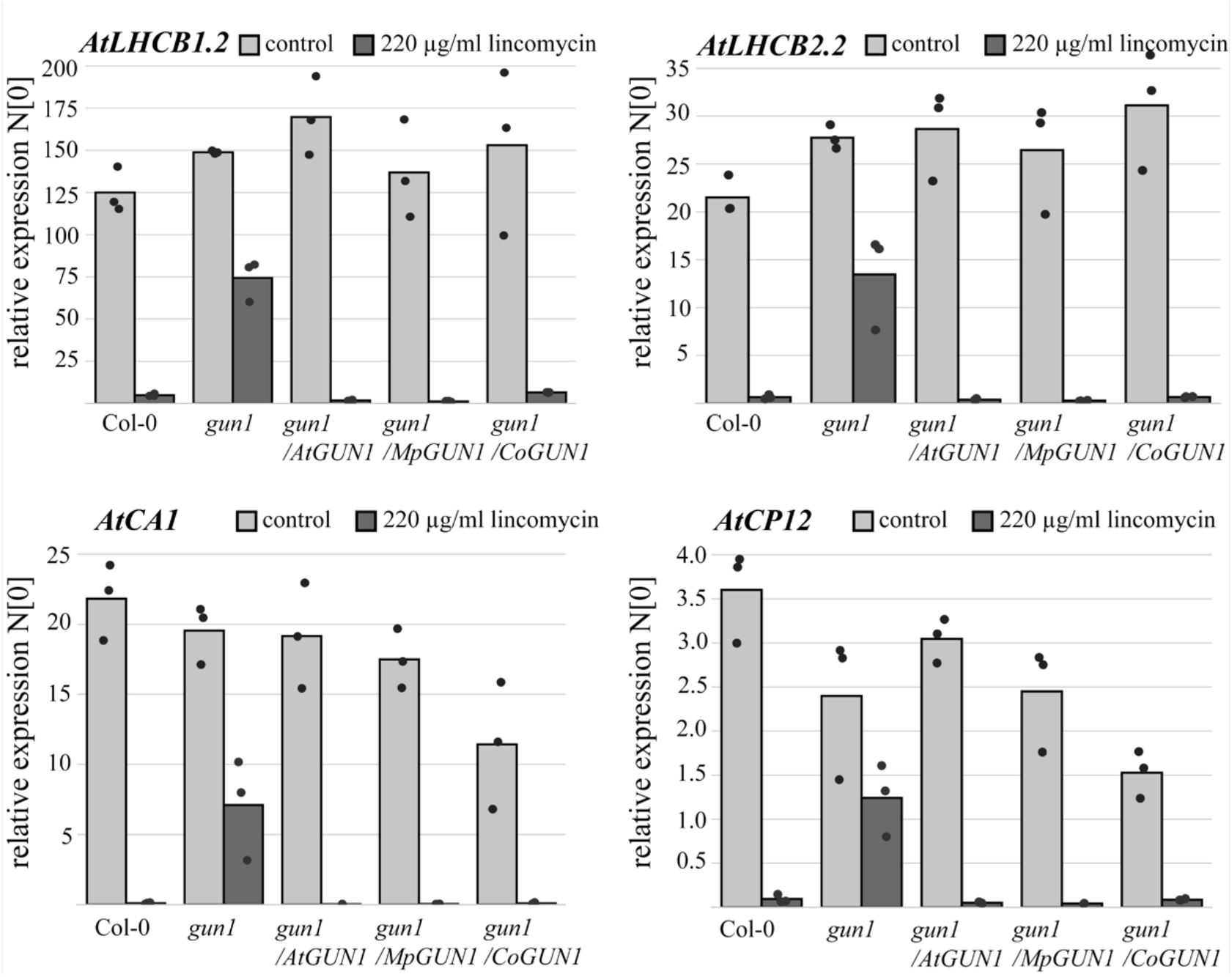
*CoGUN1* or *MpGUN1* expression in the *Atgun1* mutant restores the repression of photosynthesis-associated nuclear genes (phANG) in response to inhibition of plastid translation by lincomycin treatment. qPCR quantification of steady-state transcript levels of four phANG *AtLHCB1.2*, *AtLHCB2.2*, *AtCA1* and *AtACP12* in wild-type *Arabidopsis*, the *A*t*gun1* mutant and *Atgun1* mutant complemented with *AtGUN1* (control), *MpGUN1* or *CoGUN1*. The transcript levels were normalised against *AtPDF2* and *AtUBQ10*. Each dot represents an independent biological replicate, bars illustrate the average of the three biological replicates.

### *C. orbicularis* and *M. polymorpha GUN1* restore most aspects of the *Atgun1* mutant phenotype

As *CoGUN1* and *MpGUN1* can complement the *AtGUN1*-mediated repression of photosynthesis associated nuclear encoded genes (pHANGs) when expressed in the *Atgun1* mutant, we set out to investigate whether other aspects that involve *GUN1*-mediated signalling are also complemented.

Wild-type *Arabidopsis* seedlings germinated on media supplemented with lincomycin in the presence of 2 % sucrose develop purple cotyledons due to accumulation of anthocyanins. This anthocyanin accumulation is defective in the *Atgun1* mutant (Figure 3, Cottage *et al*., 2010). *Atgun1* seedlings complemented with *CoGUN1* or *MpGUN1* develop similar purple colouration to wild-type seedlings when germinated on lincomycin-containing media (Figure 3a). Treatment with lincomycin also inhibits cotyledon expansion in wild-type *Arabidopsis* seedlings more than in *Atgun1* mutant seedlings. *CoGUN1* and *MpGUN1* restore the inhibition of cotyledon expansion in response to inhibition of plastid translation by lincomycin (Figure 3a and 3b). These findings are consistent with the hypothesis *CoGUN1* and *MpGUN1* can activate *AtGUN1*-dependent downstream signalling that leads to upregulation of anthocyanin biosynthesis and inhibition of cotyledon expansion when plastid translation is inhibited.

**Fig. 3.**
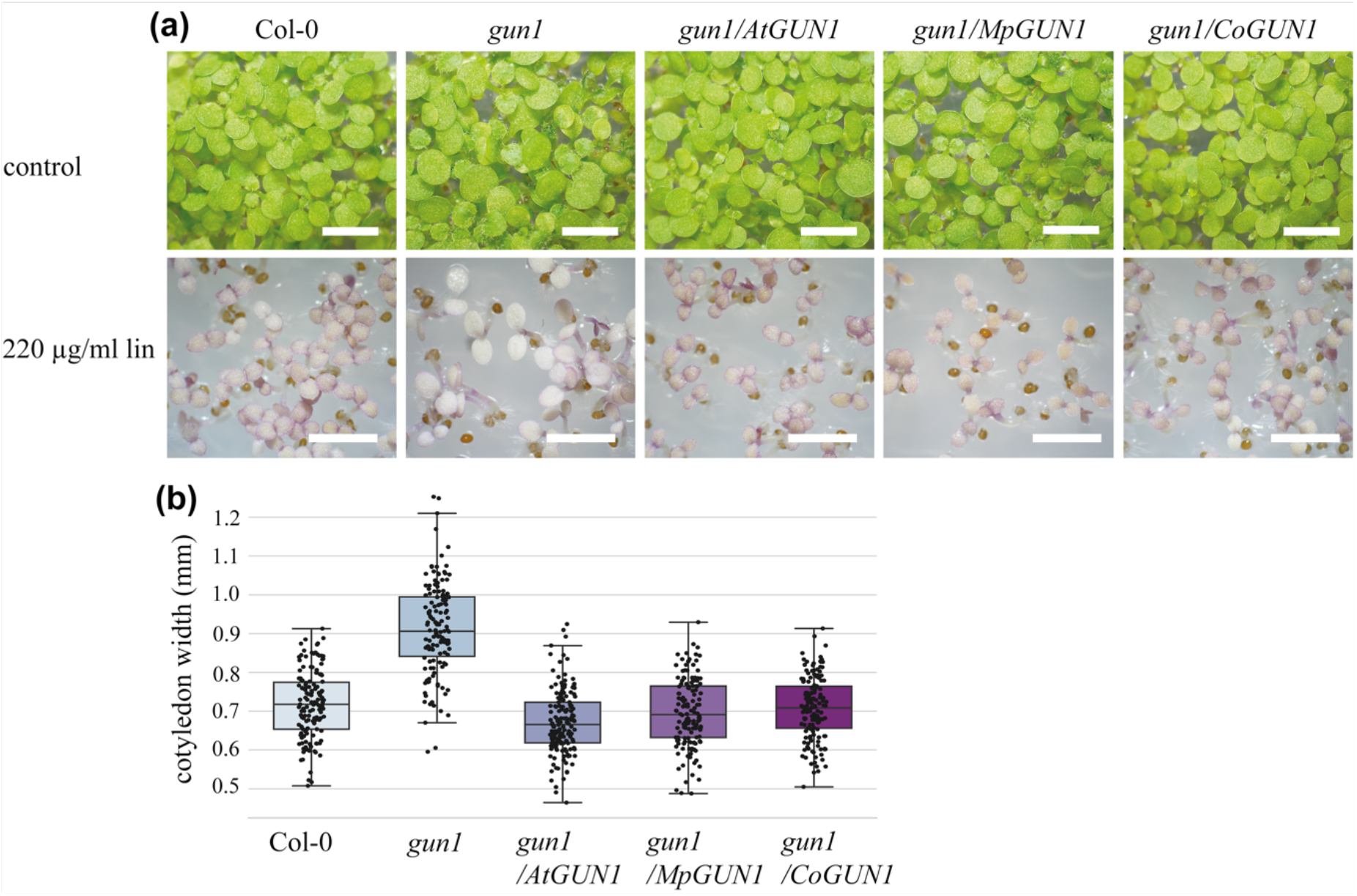
*CoGUN1* and *MpGUN1* expression in the *Atgun1* mutant restores anthocyanin accumulation and repression of cotyledon expansion in response to lincomycin treatment. a) five-day-old seedlings of wild-type *Arabidopsis*, *Atgun1* mutant and *Atgun1* mutant complemented with *AtGUN1* (control), *MpGUN1* or *CoGUN1* grown under constant light on sterile plates for 7 days (top row, scale bar 4 mm) or on media supplemented with 220 μg/ml lincomycin for 5 days (bottom row, scale bar 3 mm). b) quantification of cotyledon width of seedlings grown on media supplemented with 200 μg/ml lincomycin for 5 days. Centre line indicates the mean, box limits indicate the upper and lower quartiles, whiskers indicate the data range. Dots represent individual measurements. All other lines were significantly different from *gun1* (Tukey’s HSD: p < 0.001).

*AtGUN1* is also required for cold tolerance during germination; germinating *Atgun1* mutant seedlings are hypersensitive to cold (Marino *et al*., 2019). This cold-sensitive phenotype is barely noticeable in the cotyledons but becomes striking when the first true leaves appear after 6-7 weeks of growth at 4 °C (Figure 4). True leaves of wild-type plants germinated at 4 °C are green and expanded, while the true leaves of the *Atgun1* mutant are small, narrow and bleached. To check if *CoGUN1* and *MpGUN1* can restore the cold-sensitive phenotype of the *Atgun1* mutant we germinated the complemented seeds on agar plates at 4 °C. Both *CoGUN1* and *MpGUN1* improved the cold tolerance of the *Atgun1* mutant (Figure 4). However, *CoGUN1* only partially restored cold tolerance: the first true leaves that emerged expanded and greened more than those of the *gun1* mutant but were variegated. These findings indicate that *MpGUN1* and *CoGUN1* can at least partially complement the cold sensitive phenotype of the *Arabidopsis gun1* mutant.

**Fig. 4.**
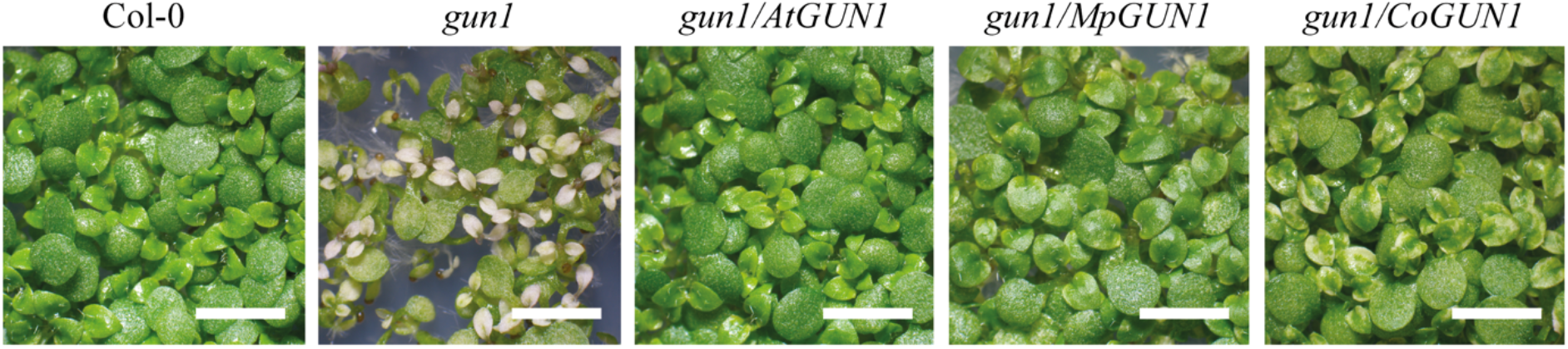
*MpGUN1* and *CoGUN1* complement the cold sensitive phenotype of the *Atgun1* mutant. Phenotype of 7-week-old seedlings germinated at 4 °C under long day light regime. Scale bar 4 mm.

*Arabidopsis* seedlings germinated in the absence of light (such as under soil) develop an etiolated phenotype characterised by an elongated hypocotyl, closed cotyledons and differentiation of proplastids into non-green etioplasts. Upon illumination (or emerging from the soil) etiolated seedlings undergo de-etiolation: hypocotyl elongation becomes inhibited, the cotyledons expand and etioplasts differentiate into photosynthetic chloroplasts. *AtGUN1*-dependent signalling operates when plastid translation is inhibited during germination under low-light conditions. Wild-type Arabidopsis seedling germinated on lincomycin-containing media under low-light conditions resemble dark-grown seedlings: their hypocotyls become elongated and cotyledons remain closed (Figure 5a-c). The *Atgun1* mutant is defective in this response, indicating that inhibition of de-etiolation in response to inhibition of plastid translation is dependent on functional GUN1 protein (Martín *et al*., 2016). *CoGUN1* and *MpGUN1* expressed in the *Atgun1* mutant background restore the inhibition of de-etiolation in response to inhibition of plastid translation by lincomycin (Figure 5). This finding suggests *CoGUN1* and *MpGUN1* can replace *AtGUN1* during de-etiolation.

**Fig. 5.**
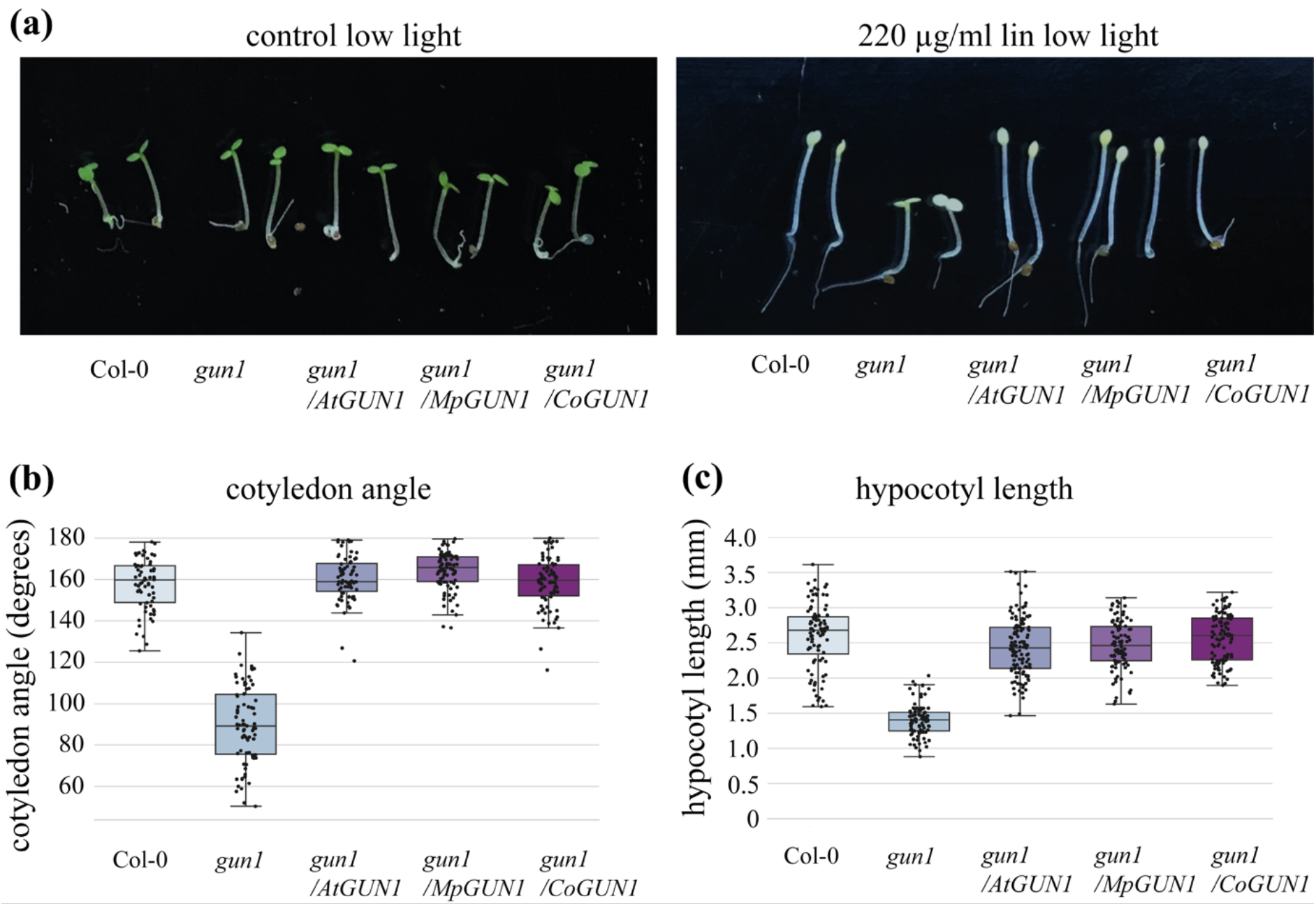
*CoGUN1* and *MpGUN1* expression in the *Atgun1* mutant restores the repression of de-etiolation when plastid translation is inhibited by lincomycin during germination. a) Wild-type Arabidopsis, *Atgun1* and *Atgun1* mutant complemented with *AtGUN1* (control), *MpGUN1* or *CoGUN1* germinated for 3 days under constant low-light (1 μmol.m^−2^.sec^−1^) on sterile plates without (control) or with 220 μg⋅ml^−1^ lincomycin. b) quantification of cotyledon angle in seedlings grown on lincomycin as in a. c) quantification of hypocotyl length in seedlings grown on lincomycin as in a. In b and c the centre line indicates the mean, box limits indicate the upper and lower quartiles, whiskers indicate the data range. Dots represent individual measurements. All other lines were significantly different from *gun1* (Tukey’s HSD: p < 0.001).

Taken together, our results indicate that GUN1 proteins from streptophyte algae, such as *C. orbicularis*, and distantly related land plants, such as *M. polymorpha*, can functionally replace the Arabidopsis GUN1 protein.

### *Mpgun1* mutants are indistinguishable from wild type under non-stressful conditions

To investigate if *GUN1* genes are functionally conserved among land plants we generated two different CRISPR deletion lines of the single-copy *GUN1* gene in the liverwort *M. polymorpha* (Figure 6a). The resulting *Mpgun1-1* and *Mpgun1-2* deletion mutant plants were PCR-genotyped, and PCR products of selected plants were sequenced to confirm the deletion introduced a premature stop codon. Male and female plants for each line were then crossed together to obtain non-segregating population of *Mpgun1-1* and *Mpgun1-2* knock-out mutant spores.

**Fig. 6.**
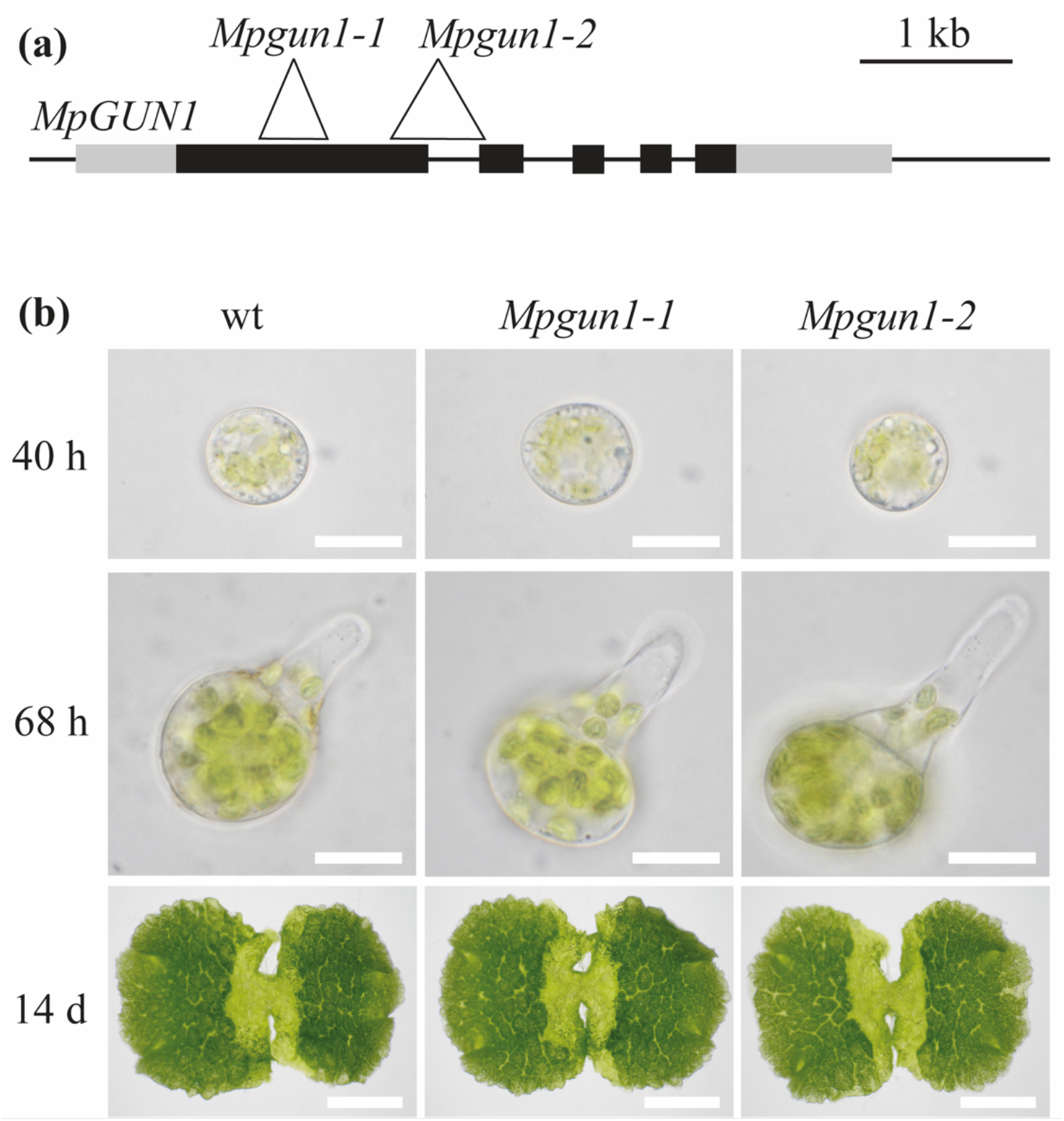
The *Mpgun1* mutant phenotype is indistinguishable from the wild-type phenotype. a) gene model showing the structure of the *M. polymorpha* GUN1 gene. Exons are indicated as black boxes, non-coding regions are indicated as grey boxes. The locations of the CRISPR-induced deletions are shown as white triangles. b) 40 hour-old (top row, scale bar 20 μm) or 68 hour-old (middle row, scale bar 20 μm) germinating spores or 14-day-old gemmae (bottom row, scale bar 3 mm) of *M. polymorpha* wild-type and *Mpgun1* mutants.

To check if *GUN1* is involved in chloroplast development in *M. polymorpha* we observed germination of wild-type and *Mpgun1* mutant spores. Wild-type *M. polymorpha* spores begin germination by cell expansion. After the first 24 hours, chlorophyll autofluorescence can be observed from the chloroplasts (Bowman *et al*., 2017). By 48 hours the spores have become fully expanded and chloroplasts appear fully developed. After 48 hours the spores undergo the first asymmetric cell division, followed by the emergence of the first rhizoid from the smaller daughter cell. Sometimes the emergence of the first rhizoid precedes the first cell division. The *Mpgun1-1* and *Mpgun1-2* knock-out mutant spores are phenotypically indistinguishable from wild type spores during spore germination (Figure 6b). This suggests *MpGUN1* is not required for chloroplast development under non-stressful conditions.

Interestingly, dark-grown *M. polymorpha* spores also develop green chloroplasts (Figure S3). This finding indicates that unlike in Arabidopsis, proplastid-to-chloroplast differentiation in *M. polymorpha* spores is independent of light signals. Consequently, proplastid-to-etioplast and etioplast-to-chloroplast transitions do not take place in *M. polymorpha* spores and therefore we conclude *MpGUN1* is not involved in these processes.

### *MpGUN1* is not involved in global repression of nuclear encoded photosynthesis-associated genes (phANGs) in response to plastid stress

The *Arabidopsis* GUN1 protein is required for chloroplast retrograde signalling and the *M. polymorpha* GUN1 protein can restore retrograde signalling in the *Arabidopsis gun1* mutant. Therefore, we hypothesised GUN1 might also be involved in chloroplast retrograde signalling in *M. polymorpha*. To test this hypothesis, we tested if germinating *Mpgun1* mutant spores are defective in chloroplast retrograde signalling when plastid translation is inhibited. *M. polymorpha* spores are almost fully resistant to lincomycin. Therefore, we used another inhibitor of plastid translation, spectinomycin, that effectively blocks *M. polymorpha* spore development (Figure S3). Spectinomycin is commonly used as a selectable marker in plastid-transformation protocols, including in *M. polymorpha* (Chiyoda *et al*., 2006; Boehm *et al*., 2016), and its mechanism of function is well-characterised (Ellis, 1970; Parker *et al*., 2014). *M. polymorpha* spores germinated on media containing 500 μg⋅ml^−1^ spectinomycin expanded and developed chloroplasts, but never grew a rhizoid or underwent the first cell division (Figure S3). This is consistent with the hypothesis that spectinomycin effectively inhibits plastid translation in *M. polymorpha.* We then checked if inhibition of plastid translation by spectinomycin during chloroplast development in *M. polymorpha* leads to reduced transcript levels of phANGs or other transcripts that are regulated by plastid retrograde signalling in seed plants. To this end, we prepared transcriptomes of wild-type and *Mpgun1* knock-out spores germinated in the absence or presence of spectinomycin. The spores were grown under long day conditions and harvested 48 hours after plating. Wild-type spores germinated on spectinomycin had reduced phANG transcript levels compared to the control (Figure 7., Table S3). However, the reduction in phANG transcript levels was much less profound than that observed in wild-type *Arabidopsis* under similar conditions and *Mpgun1* mutant spores were not defective in this response (Figure 7., Table S3). GO terms associated with photosynthesis were over-represented among the genes down-regulated in response to spectinomycin in both wild-type and *Mpgun1-1* mutant spores (Table S4). Furthermore, the set of genes that was differentially expressed in the *Mpgun1-1* mutant compared to wild-type on spectinomycin was not enriched in photosynthesis-associated GO terms, except for RbcS-encoding transcripts, which were more abundant in *Mpgun1-1* compared to wild-type (Table S4). Under control conditions the levels of phANG transcripts did not markedly differ in wild-type and *Mpgun1* mutant spores (Figure 7). These findings indicate that inhibition of plastid translation does not induce *MpGUN1-*dependent global repression of nuclear encoded photosynthesis-associated genes in *M. polymorpha*.

**Fig. 7.**
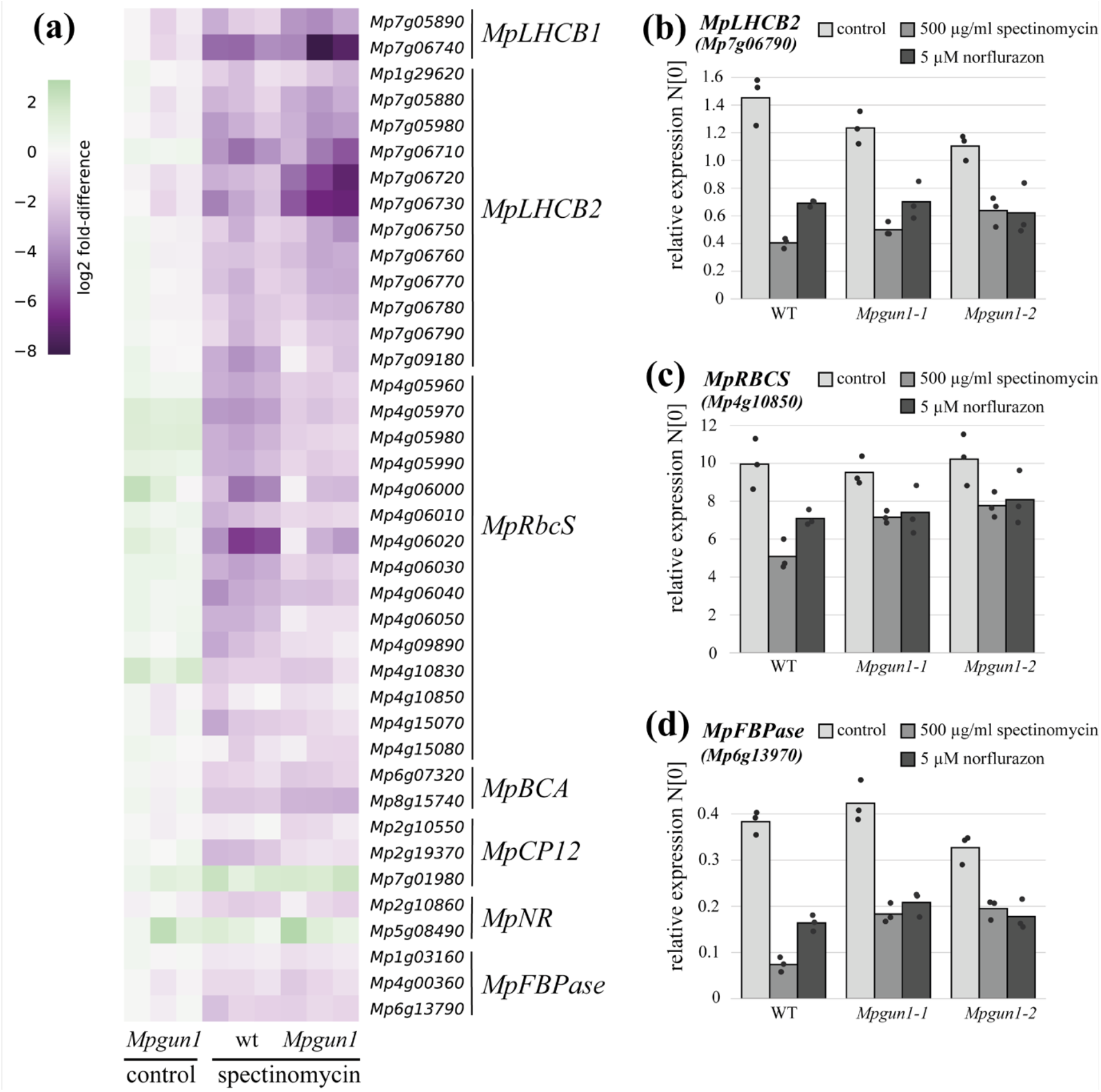
In *M. polymorpha* inhibition of chloroplast translation by spectinomycin treatment or chloroplast stress imposed by norflurazon treatment does not result in *GUN1*-mediated down-regulation of the same set of genes as in seed plants. a) Transcript levels of *M. polymorpha* orthohologs of seed plant plastid retrograde signalling-regulated genes *LHCB1*, *LHCB2, RbcS, BCA, CP12, NR and FBPase* in wild-type and *Mpgun1* mutant *M. polymorpha* spores germinated on control media or media supplemented with 500 μg⋅ml^−1^ spectinomycin. Transcript abundances are shown as log2 fold-difference compared to wild-type *M. polymorpha* spores grown under control conditions. Each column represents an independent biological replicate transcriptome. b-d) qPCR quantification of phANG transcript levels in wild-type and *Mpgun1* mutant spores germinated on spectinomycin or norflurazon-containing media. The transcript levels were normalised against *MpEF1α* and *MpACT*. Each dot represents an independent biological replicate, bars illustrate the average of the three biological replicates. The code and source data for reproducing figure 7a are obtainable from Dryad (https://doi.org/10.5061/dryad.x0k6djhmk).

To independently verify that *GUN1* is not required for inhibition of phANG expression in response to plastid stress in *M. polymorpha* we carried out qPCR quantification of phANG transcript levels wild-type and *Mpgun1* mutant spores germinated on spectinomycin or norflurazon-containing media. Norflurazon is an inhibitor of carotenoid biosynthesis. Inhibition of carotenoid biosynthesis results in oxidative stress to the chloroplasts and activates chloroplast retrograde signalling in Arabidopsis. *M. polymorpha* spores germinated on 5 μM norflurazon became bleached and their development arrested at a single cell stage (Figure S3), suggesting that norflurazon most likely also results in oxidative stress to the chloroplasts in *M. polymorpha*. We quantified the transcript levels of three phANG: *LHCB2*, RBCS and *FBPase* in spectinomycin and norflurazon treated wild-type, *Mpgun1-1,* and *Mpgun1-2* mutant spores. These phANG are present in *M. polymorpha* as multicopy genes. Therefore, primers were designed to amplify the most abundantly transcribed isoform in each gene group. Transcript levels of these phANG were reduced in spores germinated on spectinomycin or norflurazon, but *Mpgun1-1* or *Mpgun1-2* mutant spores were not defective in this response (Figure 7b-d). These findings indicate that *GUN1* is not involved in chloroplast retrograde signalling in response to plastid stress in the liverwort *M. polymorpha*.

## Discussion

GUN1 has become the most studied PPR protein since the discovery of its role in plastid retrograde signalling 15 years ago (Koussevitzky *et al*., 2007). Despite numerous attempts to pin down its function in this process, the molecular action of GUN1 remains unclear. We attempted to take a different approach by using evolutionary conservation of sequence and function to guide our interpretation of how GUN1 might act. We have shown that GUN1 is probably present in every land plant with a chloroplast genome (as well as in the streptophyte algae most closely related to land plants) and equally highly conserved in terms of its structure and sequence. Therefore, it clearly has adaptive significance to almost all plants under natural conditions. This includes many non-photosynthetic parasitic or mycoheterotrophic plants.

The question that remains is what is this essential function? Under laboratory conditions, GUN1 is entirely dispensable in *Arabidopsis* and *Marchantia*. The only conditions under which the *gun1* phenotype is truly dramatic are non-physiological treatments with inhibitors of chloroplast development such as norflurazon, spectinomycin or lincomycin, leaving its true physiological role unclear (Pogson *et al*., 2008). Nevertheless, the assumption has often been made that GUN1’s essential, conserved role is connected to retrograde signalling. For example, the first identification of *GUN1* genes in streptophyte algae led to the suggestion that the *GUN1*-mediated retrograde signalling pathway evolved prior to the colonisation of land by plants (de Vries *et al*., 2018; Nishiyama *et al*., 2018). We found that inhibition of chloroplast translation induced much less profound repression of phANGs in the liverwort *M. polymorpha* than in *Arabidopsis*. Furthermore, *Mpgun1* mutant spores were not defective in this response. This suggests that, in *Marchantia*, GUN1 is not involved in a strong global transcriptional repression of phANGs in response to plastid stress or inhibition of chloroplast translation. *M. polymorpha* spores develop green chloroplasts when germinated in complete darkness. Similarly, in gymnosperms and at least some ferns the differentiation of proplastids into chloroplasts is not light-dependent (Raghavan, 1993; Ranade *et al*., 2019). We speculate that etioplast-to-chloroplast developmental checkpoints and GUN1-mediated retrograde signalling may have only evolved in flowering plants. Our findings highlight the importance of functional characterisation of proteins in divergent model systems when inferring evolutionary conservation of signalling pathways.

It thus seems likely that the conservation of GUN1 from streptophyte algae through bryophytes, lycophytes, ferns and gymnosperms is due to its involvement in some other essential physiological process and the later involvement in retrograde signalling is a secondary role. The distribution of GUN1 across land plants and algae rules out some processes as candidates. It was recently proposed that GUN1 regulates RNA editing via directly interaction with MORF2 protein (Zhao *et al*., 2019). RNA editing has not been observed in streptophyte algae and most likely only evolved in the lineage leading to land plants (Schallenberg-Rüdinger & Knoop, 2016). Furthermore, complex thalloid (*Marchantiid)* liverworts, such as *M. polymorpha*, have entirely lost RNA editing (Groth-Malonek *et al*., 2007). Moreover, MORF2 homologues are only present in seed plants (Schallenberg-Rüdinger & Knoop, 2016; Gutmann *et al*., 2020). Therefore, the primordial role of GUN1 cannot be in regulating RNA editing. Similarly, GUN1 has been proposed to be involved in feedback regulation of protein import into plastids, but the loss of GUN1 in *Rafflesiaceae* (which still import proteins into plastids but cannot synthesise any plastid proteins) rather implies a role in chloroplast gene expression. Thus, while these recent propositions may be relevant to understanding GUN1’s secondary role in retrograde signalling, they do not appear to help identify its primary function.

Despite the fact that GUN1 is not involved in a retrograde signalling pathway in *Marchantia*, MpGUN1 fully complements an *Arabidopsis* gun1 mutant, including its retrograde signalling phenotype. This surprising result implies to us that it is not the GUN1 protein itself that directly acts in the signalling pathway, but rather the result of GUN1’s (conserved) action.

What is the primary (primordial) function of GUN1? The extreme conservation of GUN1’s PPR motifs (exceeding even that of the PPR splicing factors EMB2654 and PPR4 which have a similarly long evolutionary history, Lee *et al*., 2019b) and of its SMR domain are consistent with it functioning as a typical PPR protein, i.e. as a sequence-specific RNA binding protein. Its presence in all land plants except those lacking a plastid genome indicate that if its target is an RNA, it is one present in all plastid genomes. This rules out almost all protein-coding transcripts and tRNAs, leaving the *rrn16* and *rrn23* transcripts as the most likely targets. The extremely high conservation of the specificity-determining residues would suggest that the GUN1 binding site is also highly conserved, even in species with highly divergent plastid genomes, again consistent with a highly conserved target transcript such as an rRNA. We predict that the essential conserved role of GUN1 will turn out to be in some way involved with the regulation of plastid ribosome biogenesis.

## Acknowledgements

This work was supported by The Australian Research Council (FL140100179 and CE140100008 to IS) and CSIRO Synthetic Biology Future Science Platform (fellowship to SH). Vector pHB453 was kindly provided by Dr Holger Breuninger, University of Tübingen.

## Author Contributions

SH and IS designed the research, analysed the data and wrote the manuscript. SH performed the experiments and collected the data.

## Data Availability

Sequencing read data was deposited at the Short Read Archive database at the National Center for Biotechnology Information under project number PRJNA800059. The GUN1 HMM profile and all 893 identified GUN1 sequences aligned in FASTA format are available from Dryad (https://doi.org/10.5061/dryad.x0k6djhmk). This Dryad repository also contains the code and source data for reproducing Figures 1, 7a, S1 and S2 and Tables S2 and S3.

## Accession numbers

The following Arabidopsis genes were mentioned in this article: At*GUN1*, AT2G31400; At*LHCB1.2*, AT1G29910; At*LHCB2.2*, AT2G05070; At*CA1*, AT3G01500; and At*CP12*, AT3G62410.

The following *M. polymorpha* genes were mentioned in this article: *MpGUN1*, Mp1g08430; *MpLHCB2*, Mp7g06790; *MpRBCS*, Mp4g10850; *MpFBPase*, Mp6g13790.

## Supporting information

**Fig. S1** Multiple alignment of 76 GUN1 protein sequences from diverse streptophyte algae and land plants.

**Fig. S2** Identification of GUN1 sequences by hmmsearch scores.

**Fig. S3** Phenotype of wild-type *M. polymorpha* spores germinated under long day conditions in the absence (control) or presence of chemical inhibitors of plastid function (spec= spectinomycin 500 μg/ml or nor= norflurazon 5 μM) or in complete darkness.

**Table S1** List of primers used in this study.

**Table S2** Identification of GUN1 sequences by hmmsearch scores.

**Table S3** Differentially expressed transcripts in wild type and *Mpgun1* mutant spores grown in the presence or absence of spectinomycin.

**Table S4** Gene ontology (GO) term enrichment analysis of wild type and *Mpgun1* spores grown in the presence or absence of spectinomycin.

**Notes S1** Sequence maps of plasmids used for complementation of the *Arabidopsis gun1* mutant (Genbank format).

**Methods S1** *Agrobacterium-*mediated transformation of the liverwort *M. polymorpha.*

**Methods S2** Generation of transgenic *Mpgun1* CRISPR/Cas9 knock-out lines.

