## Supplementary figures and images for "The GENOMES UNCOUPLED1 protein has an ancient, highly conserved role in chloroplast gene expression but not in retrograde signalling"

### Fig. S1

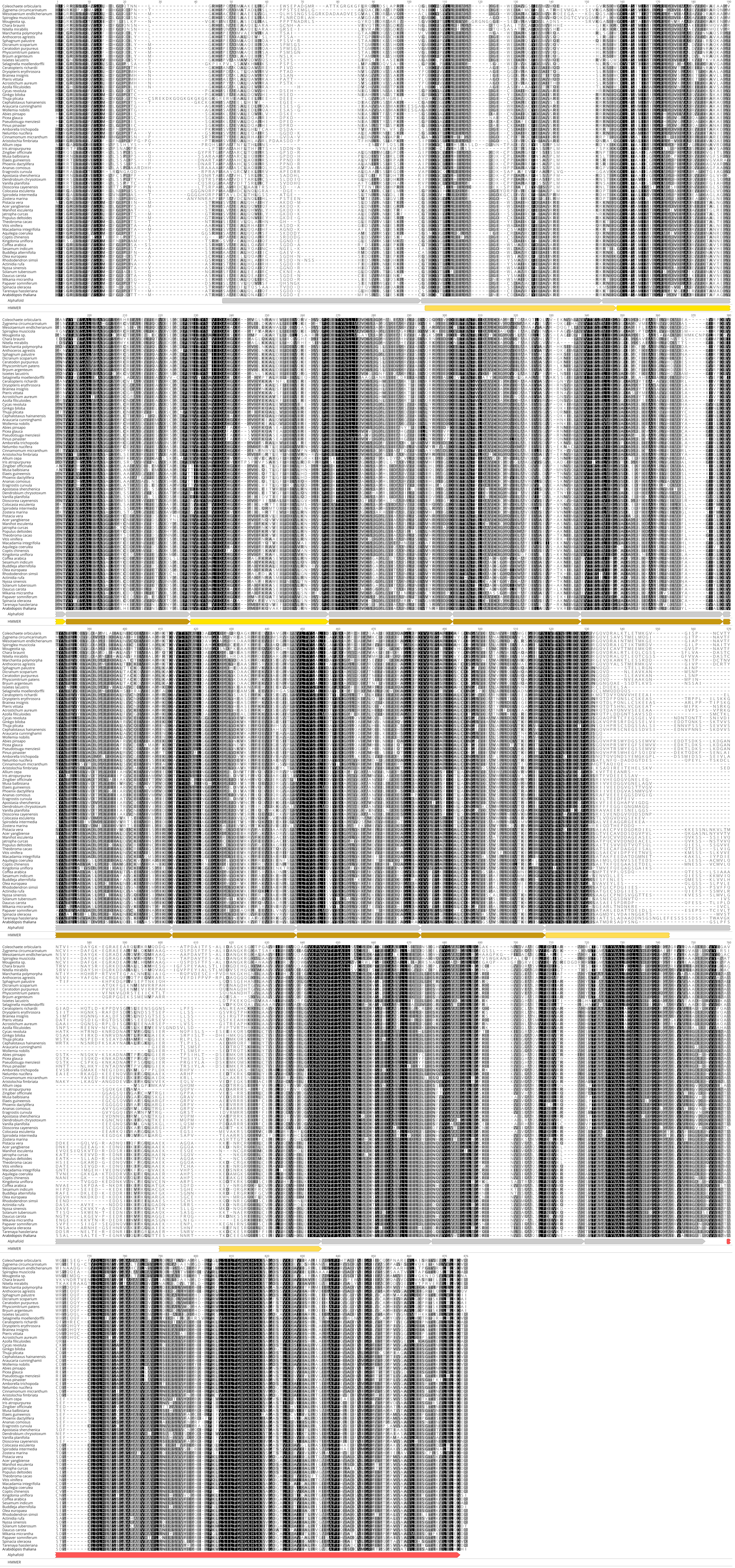

### Fig. S2

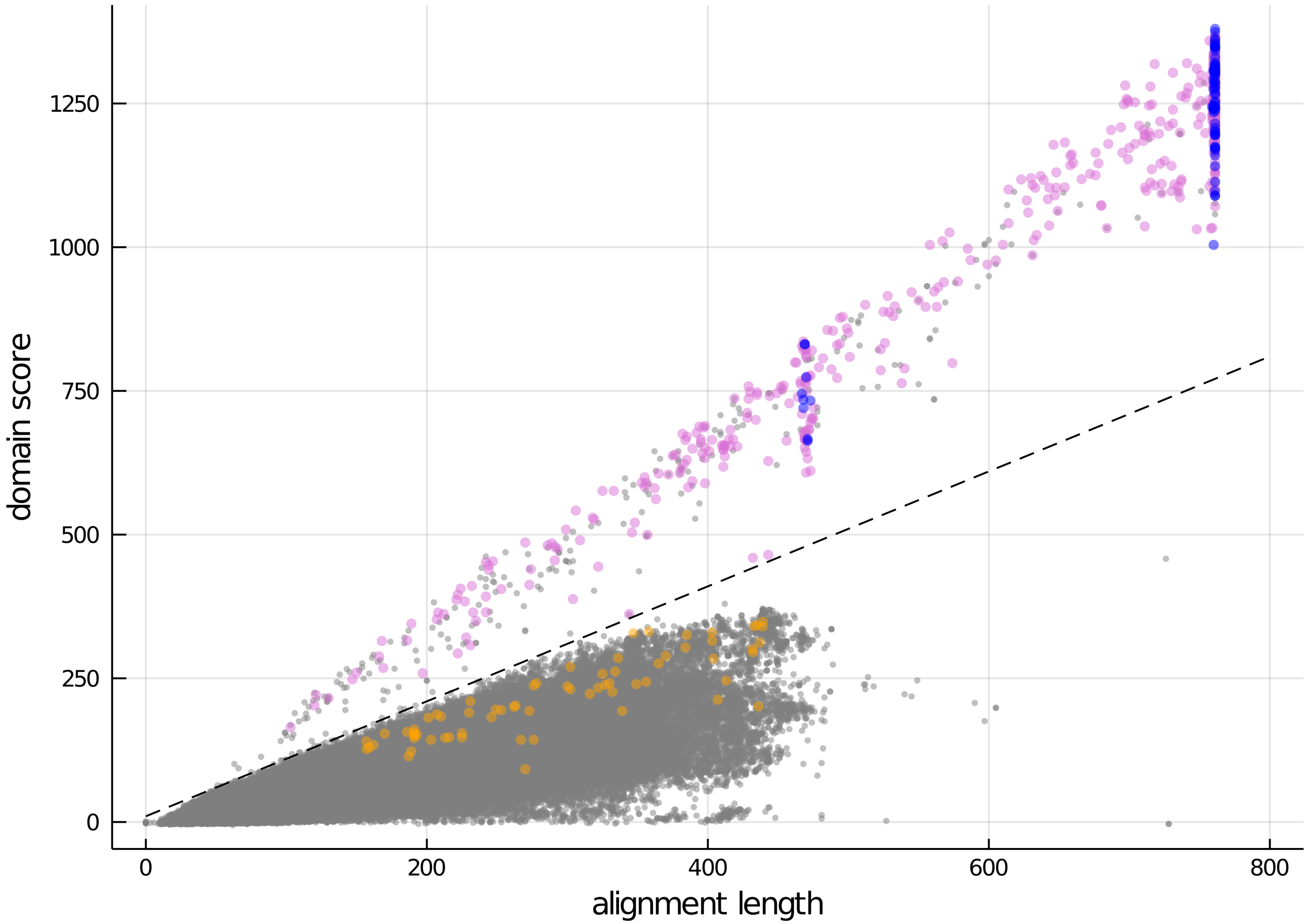

### Fig. S3

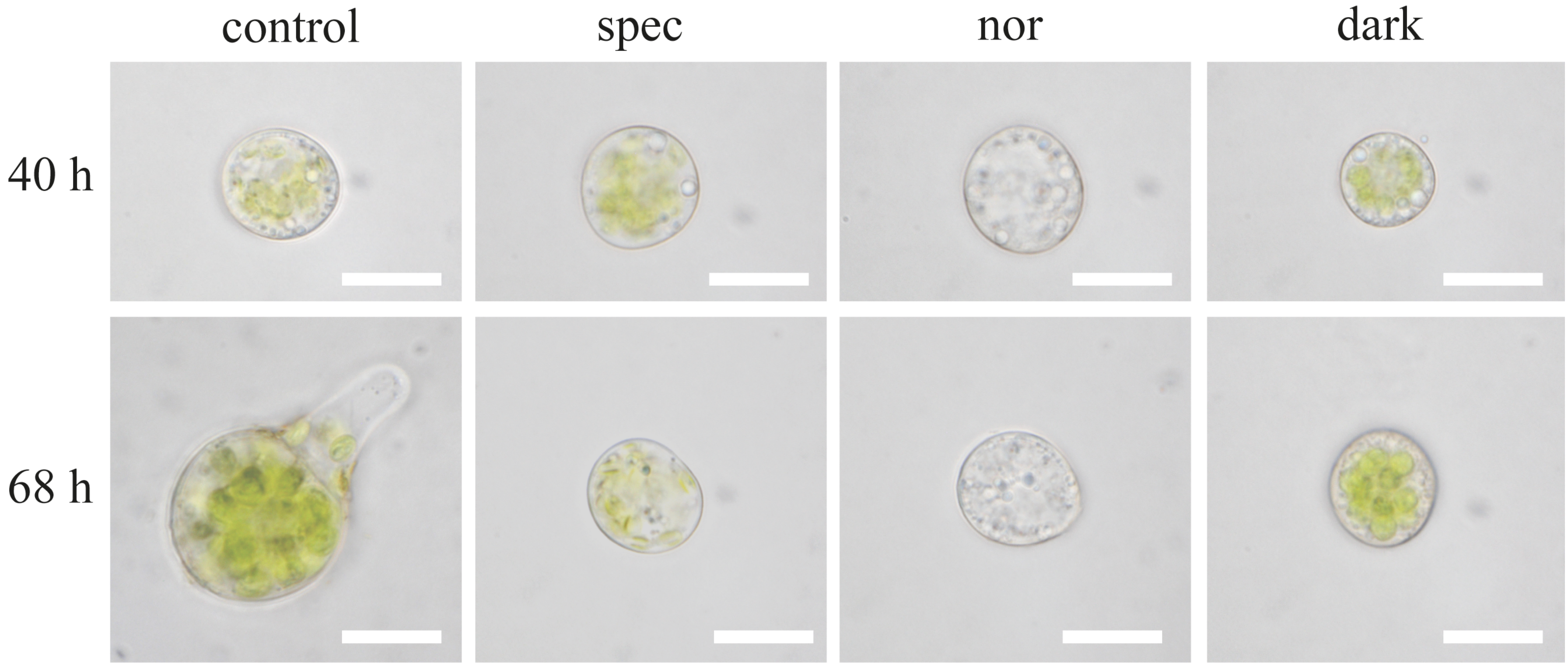
