## supporting information for "The GENOMES UNCOUPLED1 protein has an ancient, highly conserved role in chloroplast gene expression but not in retrograde signalling"

Supporting information Figures S1-S3, Tables S1-S4 and Notes S1 are provided as separate files. Supporting methods S1-S2 are provided here.

**Article title: The GENOMES UNCOUPLED1 protein has an ancient, highly conserved role in chloroplast gene expression but not in retrograde signalling**

Suvi Honkanen^1,2^, Ian Small^1*^

^1^ Australian Research Council Centre of Excellence in Plant Energy Biology, School of Molecular Sciences, The University of Western Australia, Crawley 6009, Australia

^2^ Synthetic Biology Future Science Platform, CSIRO, Canberra, ACT, Australia

**The following Supporting Information is available for this article:**

**Fig. S1** Multiple alignment of 76 GUN1 protein sequences from diverse streptophyte algae and land plants. The alignment was constructed using MAFFT and visualised with Geneious. Darker shading indicates higher similarity to the consensus. Below the alignment, the annotation tracks show helix-turn-helix motifs (grey arrows) and SMR domain (red arrow) predicted by Alphafold and PPR motifs (brown arrows) predicted by hmmsearch using a P-type PPR motif HMM. Dark brown motifs have higher confidence. The final PPR motif is interrupted by a long insertion but Alphafold predicts that the N- and C-terminal helices nevertheless interact together as in a typical PPR motif. The HMM profile and full alignment of all 893 identified GUN1 sequences are obtainable from Dryad (https://doi.org/10.5061/dryad.x0k6djhmk).

**Fig. S2** Identification of GUN1 sequences by hmmsearch scores. 554,837 PPR sequences translated from the 1KP transcriptome dataset were searched with a GUN1-specific hidden Markov model profile using hmmsearch. The maximum domain score for each protein is plotted against the alignment length. The blue points represent the score of the 76 GUN1 sequences used to construct the GUN1 profile, so can be considered as positive controls. The pink and orange points represent the highest-scoring alignment in each species in the 1KP dataset. The smaller grey points represent all other alignments, the bulk of which would not be expected to be GUN1 but rather other PPR proteins. The dashed line was drawn to separate the these assumed non-GUN1 sequences from the known GUN1 sequences; points lying above this line are assumed to represent GUN1 sequences. The code and source data for reproducing figure S2 are obtainable from Dryad (https://doi.org/10.5061/dryad.x0k6djhmk).

**Fig. S3** Phenotype of wild-type *M. polymorpha* spores germinated under long day conditions in the absence (control) or presence of chemical inhibitors of plastid function (spec= spectinomycin 500 μg/ml or nor= norflurazon 5 μM) or in complete darkness. Scale bar 20 μm.

**Table S1** List of primers used in this study.

**Table S2** Identification of GUN1 sequences by hmmsearch scores. The following columns were obtained from the 1KP data (Carpenter *et al.*, 2019; Leebens-Mack *et al.*, 2019): Species, Sample (1KP sample name), Clade, Order, Family, Tissue type (tissue from which RNA was extracted), Contamination (assessment of contamination by 1KP analysis). The following columns use data from Gutmann *et al.,* 2020: Target (ORF id number), Numprot (estimate of the number of distinct proteins encoded in the transcriptome of this sample), Tlen (target sequence length). These columns report new data from this manuscript: Fullscore (sum of scores of all alignments of this target to the GUN1 HMM), Domscore (maximum score of any alignment of this target to the GUN1 HMM), Hmmfrom (position within the HMM at which the alignment to the target begins), Hmmto (position within the HMM at which the alignment to the target stops), Alen (alignment length) and hasGUN1 (our interpretation of the scores, true if domscore > alen + 10). The code and source data for reproducing Table S2 are obtainable from Dryad (https://doi.org/10.5061/dryad.x0k6djhmk).

**Table S3** Differentially expressed transcripts in wild type and *Mpgun1* mutant spores grown in the presence or absence of spectinomycin. Differential expression analyses were carried out using DESeq2 (Love *et al.*, 2014). Functional annotations for MpTak_v5.1 genome release were used to annotate differentially expressed genes (log_2_ fold-change >1 or <-1 and padj < 0.01). The code and source data for reproducing Table S3 are obtainable from Dryad (https://doi.org/10.5061/dryad.x0k6djhmk).

**Table S4** Gene ontology (GO) term enrichment analysis of wild type and *Mpgun1* spores grown in the presence or absence of spectinomycin. Gene Ontology (GO) enrichment analyses were performed on the Dicots Plaza 4.5 platform (Van Bel *et al.*, 2018) using standard settings with differentially expressed genes showing log_2_ fold-change >1 or <-1 and padj < 0.01 as an input.

**Notes S1** Sequence maps of plasmids used for complementation of the *Arabidopsis gun1* mutant (Genbank format).

**Methods S1** *Agrobacterium-*mediated transformation of the liverwort *M. polymorpha*

**Methods S2** Generation of transgenic *Mpgun1* CRISPR/Cas9 knock-out lines.

**Methods S1** ***Agrobacterium-*mediated transformation of the liverwort** ***M. polymorpha***

This protocol is described in more detail in Honkanen and Jones, 2020. To obtain *M. polymorpha* spores male and female plants were grown on soil at 22 °C under 16 hours light: 8 hours dark photoperiod supplemented with far-red light and crossed as described in (Chiyoda *et al.*, 2008). 2-7 intact or burst mature sporangiums were collected in each Eppendorf tube and the opening of the tube covered with a Sun Cap Closure 18 mm (Sigma) or micropore tape. To dry the spores the tubes were first placed lids open in an air-tight container containing silica gel for 10 days and then stored at -80 **°**C. Before use the spores were briefly defrosted at room temperature and sterilised in 0.1 % sodium dichloroisocyanurate (Sigma) solution for 5 minutes. The spores were collected by centrifugation at 13,000 rpm for 1.5 minutes, after which the sterilisation solution was discarded, and spores resuspended in distilled H_2_O (100 μl per sporangium). Liquid spore cultures were prepared in 6 well plates (Sarstedt) using 6 ml sterile ½ Gamborgs medium (for 500 ml media: 0.8 g ½ Gamborgs medium powder (Duchefa Biochemie), 10 g sucrose, 150 ml L-glutamine (Sigma), 250 mg MES buffer (Sigma), pH to 5.6 using KOH) in each well. 100 μl sterilised spore solution was placed in each well and the plate was sealed with micropore tape. Spores were grown in a growth cabinet for 7 days at 22 °C constant light at 80 µE.m-2 .s-1 without shaking. Agrobacterium GV3101 liquid cultures transformed with the relevant plant transformation vectors were inoculated from single colonies into 5 ml LB medium cultures containing the relevant antibiotics and grown for 2 days at 28 °C 200 rpm on a shaker. The bacterial cultures were centrifuged 15 minutes at 2000 g, after which the supernatant was discarded and bacterial pellet resuspended in 10 ml fresh ½ Gamborgs medium containing 100 uM acetosyringone. The Agrobacterium cultures were induced by growing for 4 hours at 28 °C 200 rpm on a shaker. 100 μl of induced bacterial culture was added into each 6 ml spore culture and acetosyringone was added to final concentration of 100 uM. Spores were co-cultivated with agrobacterium for 1-3 days at 22 °C under 16 hours light 8 hours dark photoperiod or under constant light at 80 µE.m-2 .s-1 on a 120 rpm shaker. To remove the agrobacteria the spore culture was collected using a sterile plastic Pasteur pipette (Thermo Fisher), transferred onto a 40 µm nylon mesh cell strainer (Fisher Scientific) placed in the opening of a 50 ml falcon tube and rinsed with 50 ml sterile distilled H_2_O. The spores were then plated on plates containing sterile ½ Gamborgs medium pH 5.6 supplemented with 1 % (w/v) sucrose and 1.4 % (w/v) agar (Sigma) and appropriate antibiotics (always 100 ug/ml Cef to kill the agrobacterium, in addition 10ug/ml Hyg or 0.5 uM chlorosulfron to select for transformant plants). Plates were placed in a growth chamber at 22 °C, under constant light or 16 hours light 8 hours dark photoperiod. After 1-2 weeks antibiotic resistant transformant plants were visible.

**Methods S2 Generation of transgenic *Mpgun1* CRISPR/Cas9 knock-out lines**

*M. polymorpha* CRISPR/Cas9 knock-out lines were generated as described in (Sugano *et al.*, 2018). A new sgRNA construct pHB453 that allows the use of two sgRNAs that specify two adjacent CRISPR/Cas9 cut sites was kindly provided by Dr Holger Breuninger, University of Tübingen. The design of pHB453 is based on pMpGE_En03 (Sugano *et al.*, 2018, Addgene plasmid #71535), with a second propU6-sgRNA fragment introduced between the *att* sites. The first sgRNA was introduced into the *BpiI* site of pHB453 using primers MpGUN1_CRISPR_1-1 F and R for the *Mpgun1-1* construct and MpGUN1_CRISPR_2-1 F and R for the *Mpgun1-2* construct. The second sgRNA was introduced into the *BsmBI* site of pHB453 using primers MpGUN1_CRISPR_1-2 F and R for the *Mpgun1-1* construct and MpGUN1_CRISPR_2-2 F and R for the *Mpgun1-2* construct. The resulting entry vectors were sequence-verified by Sanger sequencing, after which the plasmids were recombined with the destination vector pMpGE011 (Sugano *et al.*, 2018) using Gateway LR clonase (Invitrogen).

*M. polymorpha* transformation with the binary vectors described above was carried out using *Agrobacterium* (strain GV3101) as described in Methods S1. Genotyping of *Mpgun1-1* and *Mpgun1-2* CRISPR knock-out lines was carried out using the Phire plant direct PCR kit (Thermo Fisher) as recommended by the manufacturer with primers amplifying across the predicted deletion site (MpGUN1_F1 and MpGUN1_R1 for *Mpgun1-1*, MpGUN1_F2 and MpGUN1_R2 for *Mpgun1-2*). The positive lines identified were propagated through gemmae and re-genotyped to obtain stable non-chimeric knock-out lines. The genotyping PCR products were Sanger-sequenced at Macrogen, South Korea to identify lines where the CRISPR deletion resulted in a premature stop codon. Selected T0 plants for each mutant line (*Mpgun1-1* or *Mpgun1-2*) were crossed together to obtain a non-segregating T1 population of knock-out mutant spores. T1 or T2 spores were used for phenotype assessment and gene expression analyses. The spores were re-genotyped after each experiment to confirm the absence of wild-type *GUN1* DNA.

**Honkanen, S., Jones V. A. S. 2020**. A simplified protocol for *Agrobacterium*-mediated transformation of the liverwort *Marchantia polymorpha*. *protocols.io* https://dx.doi.org/10.17504/protocols.io.ba28ighw

**Sugano SS**, **Nishihama R**, **Shirakawa M**, **Takagi J**, **Matsuda Y**, **Ishida S**, **Shimada T**, **Hara-Nishimura I**, **Osakabe K**, **Kohchi T**. **2018**. Efficient CRISPR/Cas9-based genome editing and its application to conditional genetic analysis in *Marchantia polymorpha*. *PloS one* **13**: e0205117-22.

**Chiyoda S**, **Ishizaki K**, **Kataoka H**, **Yamato KT**, **Kohchi T**. **2008**. Direct transformation of the liverwort Marchantia polymorpha L. by particle bombardment using immature thalli developing from spores. *Plant cell reports* **27**: 1467–1473.

**Sugano SS**, **Nishihama R**, **Shirakawa M**, **Takagi J**, **Matsuda Y**, **Ishida S**, **Shimada T**, **Hara-Nishimura I**, **Osakabe K**, **Kohchi T**. **2018**. Efficient CRISPR/Cas9-based genome editing and its application to conditional genetic analysis in Marchantia polymorpha. *PloS one* **13**: e0205117-22.

**Wheeler TJ**, **Clements J**, **Finn RD**. **2014**. Skylign: a tool for creating informative, interactive logos representing sequence alignments and profile hidden Markov models. *BMC bioinformatics* **15**: 7.
